# Diurnal and seasonal regulation of brain mu-opioid and dopamine D2 receptor signalling in humans

**DOI:** 10.64898/2026.09.01.748534

**Authors:** Michal Rafal Zareba, Juha Rinne, Jarmo Hietala, Pirjo Nuutila, Lauri Nummenmaa, Lihua Sun

## Abstract

**Purpose:** Rhythmic regulation of human neurotransmission systems, particularly the mu-opioid receptor (MOR) and dopamine D2 receptor (D2R) signaling, remains poorly understood. Although prior studies indicate possible diurnal fluctuations in D2R signaling, potential existence of diurnal rhythms in MOR signaling has yet to be investigated. Furthermore, whether and how seasonal factors may override or interact with diurnal rhythms in either system remains unexplored. Decoding these temporal dynamics and their possible links to behavioral traits, such as anxiety, may improve mood regulation models and inspire rhythm-tailored interventions.

**Methods:** Based on a historical PET database of healthy humans scanned with [^11^C]carfentanil (n = 188) and [^11^C]raclopride (n = 184), we investigated the diurnal patterns of MOR and D2R availability in the brain, using time-of-day (TOD) of scanning as predictor. Furthermore, we examined whether the TOD differences in receptor availability were modulated by seasonal differences in the daylength (D2R and MOR) and trait-anxiety (MOR).

**Results:** The data revealed increased MOR and decreased D2R availability in the afternoon compared to the morning scans. Interaction with the seasonal effect was, however, observed only for MOR availability, spanning brain areas including anterior cingulate, dorsomedial and dorsolateral prefrontal cortex, insula and striatum. Besides, the diurnal variation of MOR availability in these regions was also sensitive to trait-anxiety levels.

**Conclusions:** The current results reveal the diurnal patterns in opioidergic and dopaminergic neurotransmission with an opposing trajectory. Furthermore, the data highlight a unique role of the MOR system, mediating both types of rhythmic regulations, as well as behavioral traits.

## 1. Introduction

In the course of evolution, multiple species, including humans, have adapted their functioning to the naturally occurring cycles of light and dark, exhibiting diurnal and seasonal rhythms [Cajochen and Schmidt, 2025]. Adaptive physiological changes are important for survival and well-being, and these functional alterations involve the neurotransmitter signaling pathways crucial for mood changes and behavior [Zhang and Volkow, 2023; Sun et al., 2021; Sun et al., 2024].

As an immediate consequence of light-dark cycles, the periods of wakefulness and sleep, together with the associated changes in physiological parameters, such as core body temperature or cortisol and melatonin levels, repeat themselves within 24-hour cycles. Diurnal fluctuations are observed in objective measures of cognition, particularly in the attentional and executive domains, and subjective measures of mood, motivation and effort [Lo et al., 2013; Santhi et al., 2016; Burke et al., 2015; Balter et al., 2024]. Functional magnetic resonance imaging (fMRI) studies have demonstrated that these behavioral characteristics are reflected at the level of brain activity. For instance, time-of-day (TOD) modulates striatal responses to monetary rewards [Byrne et al., 2017a] and the activity of the attentional networks [Marek et al., 2010], with the lowest activation levels observed in the afternoon. In the meanwhile, reactivity of amygdala to threat-related stimuli decreases linearly with TOD [Baranger et al., 2017]. These findings suggest a potential variability in diurnal fluctuations observed across distinct neural systems.

Nonetheless, the neuromolecular mechanisms underlying these functional changes remain largely unknown. A seminal study on the serotonergic system revealed diurnal increase in the availability of cortical 5-HT_1A_ receptors and a decrease in the availability of 5-HTT transporter in the midbrain [Matheson et al., 2015]. These markers of serotonergic neurotransmission strength might be, however, relevant mainly for understanding diurnal variability in processes associated with negative reinforcement [Yee et al., 2021]. Potential neurotransmission systems implicated in the wider range of described diurnal phenomena include the mu-opioidergic and dopaminergic signaling pathways. Mu-opioid receptors (MOR) contribute to processing of both positive and negative emotions, as well as pain [DaSilva et al., 2019; Jern et al., 2023; Seppälä et al., 2026; Zubieta et al., 2003; Hsu et al., 2013; Sun et al., 2022]. Dopamine is similarly known for processing of positive and negative affective information [Hahn et al., 2021], possibly reflecting its motivational value [Berridge and Robinson, 2016]. Moreover, it modulates attentional and executive functioning [Volkow et al., 2009; Monchi et al., 2006].

Several lines of evidence suggest that MOR and dopaminergic signaling may vary in the course of a day. The plasma levels of β-endorphins, one of the endogenous MOR agonists, decrease in humans from the morning to the afternoon [Hindmarsh et al., 1989], with opioid labor analgesia requests following the opposite trajectory [Scavone et al., 2010]. Furthermore, the number of lethal opioid overdoses and the necessary dose of naloxone for revival also peaks at the late night and early morning [Gallerani et al., 2001], together suggesting highest endogenous MOR signaling levels around the beginning of the active phase in humans (i.e. early morning). With regards to dopamine, one study reported no significant differences in striatal dopamine type-2 receptor (D2R) availability between morning and late evening in humans [Earley et al., 2013]. However, a more recent work in animals demonstrated that the tonic dopamine levels were the highest around the middle of the active phase (i.e. afternoon for humans), with comparable concentrations observed at the beginning and end of the wakefulness [Ferris et al., 2014]. The possibility of observing a similar curvilinear variability of dopaminergic neurotransmission in humans, as well as the presence of TOD-related differences in central MOR signaling, is nonetheless yet to be validated.

Physiological processes operate across multiple timescales, from diurnal rhythms aligned with the daily light–dark cycle to infradian rhythms occurring over longer periods, including the seasonal rhythms shaped by annual variations in daylength. Our recent work has demonstrated seasonal variation in both MOR and dopamine signaling in humans and in experimental rats [Sun et al., 2021, 2022, 2024, 2025]. However, it remains unclear how these seasonal patterns interact with diurnal variation in the same systems. Examining rhythmic variation across both timescales may provide a more comprehensive understanding of their physiological regulation and establish a baseline for identifying potential pathological alterations.

Therefore, in the current work we examined the diurnal fluctuations in the MOR and D2R systems in the human brain, investigating whether these patterns were further modulated by their seasonal rhythms. Our previous report has also linked trait-anxiety with alterations in the MOR system [Nummenmaa et al., 2020]. As such, we additionally tested whether trait-anxiety modulated specifically the diurnal MOR trajectories. We hypothesized that central MOR availability would increase later in the day, consistent with previously reported reductions in endogenous opioid levels. We further expected that the seasonal factors and affective traits may either mask or amplify this diurnal pattern. Last but not least, we explored potential diurnal patterns in the D2R system without a priori hypotheses.

## 2. Materials and methods

### 2.1. Data

The study was based on a retrospective human positron emission tomography (PET) data retrieved from the AIVO database (https://aivo.utu.fi) of the Turku PET Center. Baseline scans from individuals with no neurological or psychiatric conditions were used. MOR availability was examined with [^11^C]carfentanil PET (n = 188, 63.83% males; mean age = 32.94 ± 10.39 years). D2R availability was measured with [^11^C]raclopride PET (n = 184, 79.35% males; mean age = 31.68 ± 13.09 years). Individuals were scanned only once for an examination.

To investigate the diurnal effects, the TOD of scanning (hours) was used as the predictor. The TOD values ranged from 9:03 to 17:19, and linear diurnal effects were tested. To assess seasonality impact, we calculated the daylength on the day when a PET image was collected, as in our previous studies [Sun et al., 2021; Sun et al., 2024]. For a subset scans with [^11^C]carfentanil (n = 100, 78% males; mean age = 31.26 ± 11.02 years), additional measurements of trait-anxiety were acquired with the State-Trait Anxiety Inventory (STAI-X) [Spielberger, 1989].

### 2.2. PET image analysis

PET data was preprocessed using the Magia toolbox (https://github.com/tkkarjal/magia) [Karjalainen et al., 2020]. This consisted of motion correction and coregistration of PET and anatomical MRI data. The full-volume parametric PET data were obtained by normalising individual data to the MNI template and smoothing it with a 8 mm Gaussian filter. Tracer binding was quantified as BP_ND_, i.e. the ratio of specific binding to nondisplaceable binding of radioligand, using the simplified reference tissue model (SRTM) [Lammertsma and Hume, 1996]. The reference regions were delineated in the native space using FreeSurfer (https://surfer.nmr.mgh.harvard.edu/), and were located in the occipital cortex and cerebellar grey matter for MOR and D2R data, respectively.

For the MOR data, both full-volume and region-of-interest (ROI) analysis was performed. Analysis for the D2R images, given the low reliability of cortical [^11^C]raclopride binding [Farde et al., 1985], was restricted to the striatal ROIs. The lateral hemispheric structures were delineated using individual FreeSurfer-based parcellations: caudate nucleus, nucleus accumbens, pallidum and putamen.

### 2.3. Analysis of MOR availability data

The full-volume MOR availability data was analysed using AFNI’s *3dMVM* program [Chen et al., 2014]. Three types of regression models were examined. Firstly, using the full sample (n = 188), the analysis focused on the main effects of diurnal rhythmicity (i.e. TOD) and its interactions with daylength. As indicated by previous findings [Sun et al., 2021), daylength was included in the models as the second-order polynomial, together with its linear component. Age, sex and scanner type were controlled for in the same model.

Two additional models were run using the subsample that has trait-anxiety measures (n = 100). In the first model, we aimed to derive information on how diurnal variability in MOR availability was linked with individual anxiety levels. The model included both main effects and interactions of TOD with trait-anxiety, while seasonality effects, age, sex and scanner types were included as nuisance regressors. To shed light on the potential interplay between trait-anxiety and seasonal factors, another model was used to test their main effects and interaction effect, keeping TOD, sex, age and scanner type as covariates. All the findings were corrected for multiple comparisons at the cluster level using the family-wise error rate method (p_FWE_ < 0.05).

Beside the full-volume analysis, we performed ROI analysis deploying the linear mixed effect models in the *lme4* package in R (version 4.2.1). Using the Automated Anatomical Labelling 3 (AAL3) atlas [Rolls et al., 2019], we selected 28 areas known for their contributions to affective processing in humans [Nummenmaa and Tuominen, 2018; Sun et al., 2021] (see **Supplementary Tables S1 and S2**). ROI-wise values were extracted from unsmoothed normalised BP_ND_ images, and were log-transformed. In the analyses, we used analogous models to the whole-brain calculations, with the exception of using scanner type as the random factor. The results were corrected for multiple comparisons using the false discovery date (FDR) method (p_FDR_ < 0.05).

Given that the associations of daylength or subclinical anxiety with MOR availability have been reported [Sun et al., 2021; Nummenmaa et al., 2020], the current study will focus on the main effects of TOD, its interactions with trait-anxiety and daylength, as well as the interactions between trait-anxiety and daylength.

### 2.4. Analysis of D2R availability data

The regional D2R data was analysed with the linear mixed effect models available using the *lme4* package in R (version 4.2.1). Data were log-transformed, and modelled separately for each ROI using the main effects and interaction effect of TOD and daylength, keeping age and sex as covariates, and using scanner type as the random intercept. The results were corrected for multiple comparisons with the FDR method. As the main effects associated with seasonality have been reported [Sun et al., 2024], in the current work we will describe only the results pertaining to the main effects of TOD and its interactions with daylength.

## 3. Results

### 3.1. Analysis of MOR availability data

The full volume analysis of MOR availability data revealed a positive association between the MOR availability and TOD (cluster-level p_FWE_ < 0.001; **Figure 1A**). The identified regions spanned bilateral medial frontal and dorsolateral prefrontal cortex, medial temporal lobes and putamen, as well as the left postcentral gyri and lateral temporal cortex. Findings of the ROI analysis went in line with those of the full-volume data (**Supplementary Table S1**). **Figure 1B** shows morning-to-afternoon increase of MOR receptor availability in selected ROIs.

**Figure 1.**
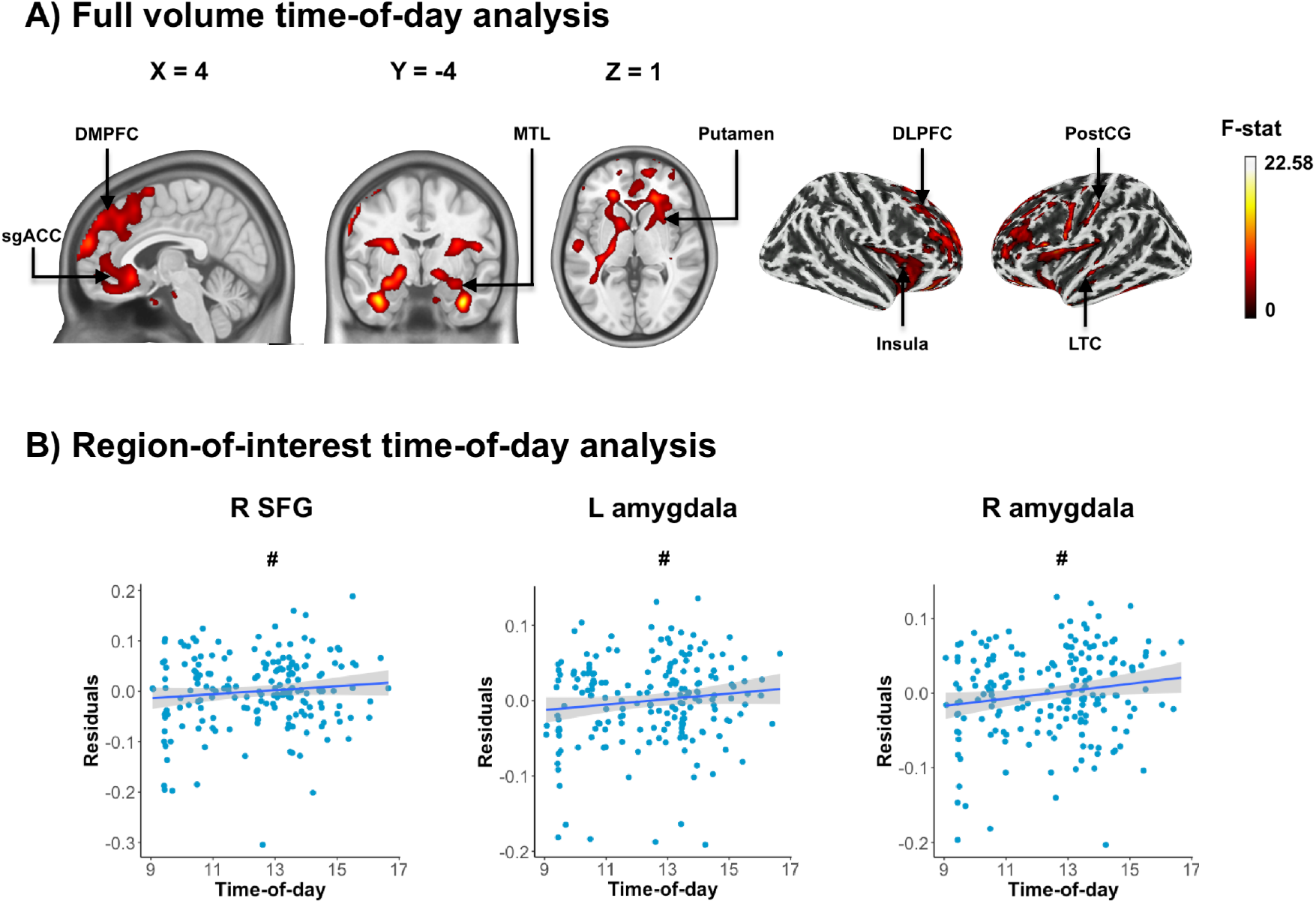
Human brain mu-opioid receptor (MOR) availability is associated with time-of-day (TOD). **A)** The results of the full volume analysis (cluster p_FWE_ < 0.001). **B)** Diurnal effects in selected regions-of-interest (ROIs). TOD was plotted against the residuals from the statistical models that excluded the main effect of TOD. Hash (#) indicates that the ROI-level findings reached nominal significance level (p_uncorr._ < 0.05). Full ROI data are found in the Supplementary Table S1. Abbreviations: L, left; R, right; sgACC, subgenual anterior cingulate cortex; DMPFC, dorsomedial prefrontal cortex; MTL, medial temporal lobe; DLPFC, dorsolateral prefrontal cortex; PostCG, postcentral gyrus; LTC, lateral temporal cortex; SFG, superior frontal gyrus.

The full volume data also showed significant interaction between TOD and linear daylength term, and between TOD and trait-anxiety (both cluster-level p_FWE_ < 0.001). The interactions with daylength were observed in bilateral dorsolateral and medial frontal regions, extending to the more posterior cortical midline structures, including paracentral lobule, posterior cingulate, precuneus and cuneus (**Figure 2A**). These associations were also present in large parietal and temporal areas, insula, thalamus and striatum.

**Figure 2.**
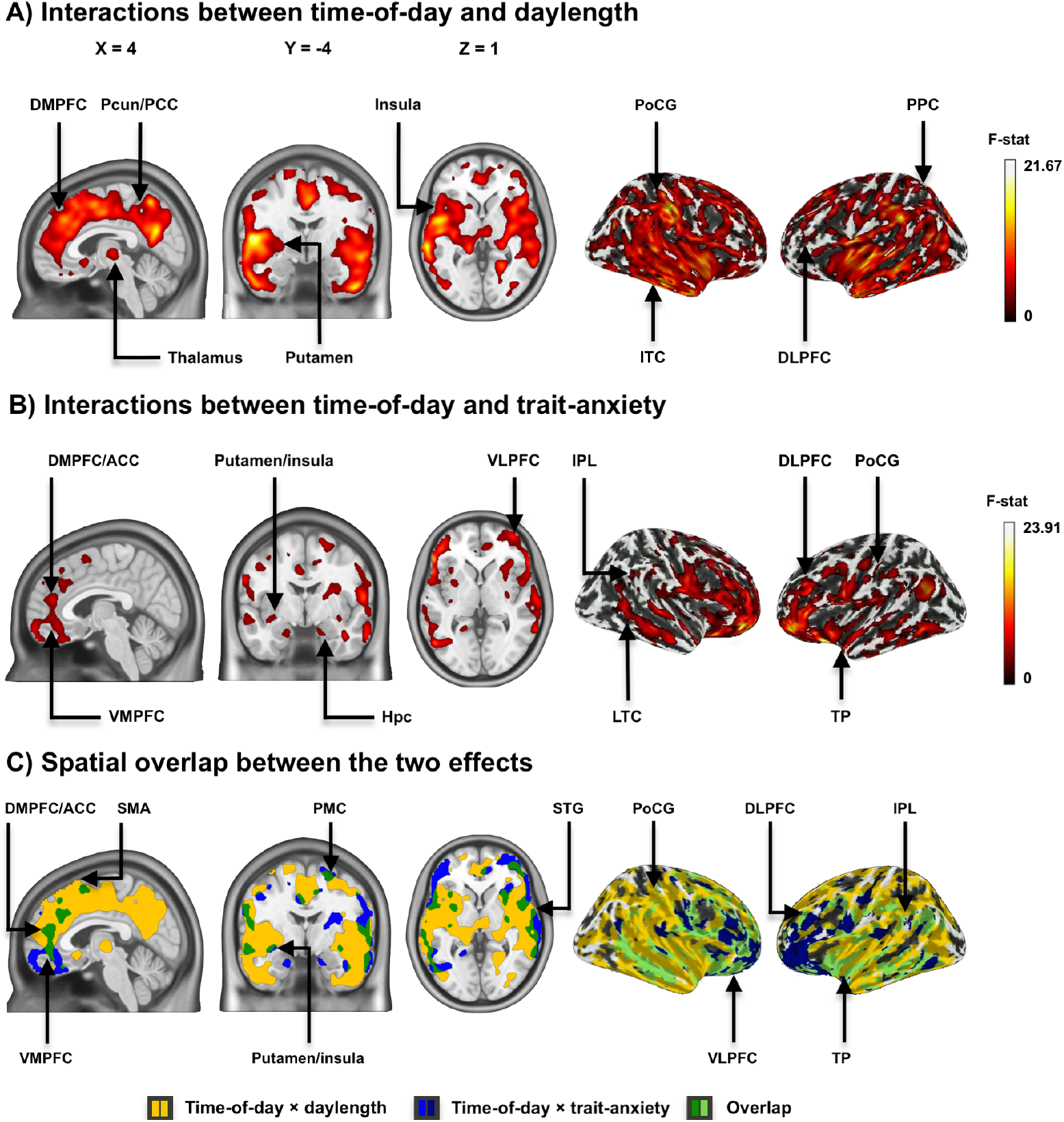
Mu-opioid receptor (MOR) availability is sensitive to the interaction effects of time-of-day with daylength (**A**) and the trait-anxiety levels (**B**). Findings were thresholded at p < 0.05, and family-wise error rate (FWE)-corrected at the cluster level. The two interaction effects were observed in overlapping brain regions (**C**). Abbreviations: DMPFC, dorsomedial prefrontal cortex; Pcun, precuneus; PCC, posterior cingulate cortex; PoCG, postcentral gyrus; ITC, inferior temporal cortex; DLPFC, dorsolateral prefrontal cortex; PPC, posterior parietal cortex; ACC, anterior cingulate cortex; VMPFC, ventromedial prefrontal cortex; Hpc, hippocampus; VLPFC, ventrolateral prefrontal cortex; IPL, inferior parietal lobule; LTC, lateral temporal cortex; TP, temporal pole; SMA, supplementary motor area; PMC, premotor cortex; STG, superior temporal gyrus.

In contrast, regions that showed interaction effects of TOD and trait-anxiety spanned bilateral medial, ventrolateral and dorsolateral frontal regions, polar and lateral temporal areas, inferior parietal lobules, hippocampal formation, insula and putamen, as well as the right postcentral gyrus (**Figure 2B**). The two types of interaction effects overlapped in bilateral frontal regions, including anterior cingulate, orbitofrontal, dorsomedial and dorsolateral prefrontal cortex, as well as in insula, putamen, inferior parietal lobe, temporal areas and the right postcentral gyrus (**Figure 2C)**.

Schematic plotting of the data shows that the MOR availability increased with TOD predominantly under longer daylength (**Figure 3A**). Similarly, this pattern of TOD-related differences was more visible in participants with lower anxiety levels (**Figure 3B**). Shorter daylength and higher anxiety levels tended to diminish these effects. The ROI analyses validated the findings of the full-volume data (**Supplementary Tables S1 and S2**).

**Figure 3.**
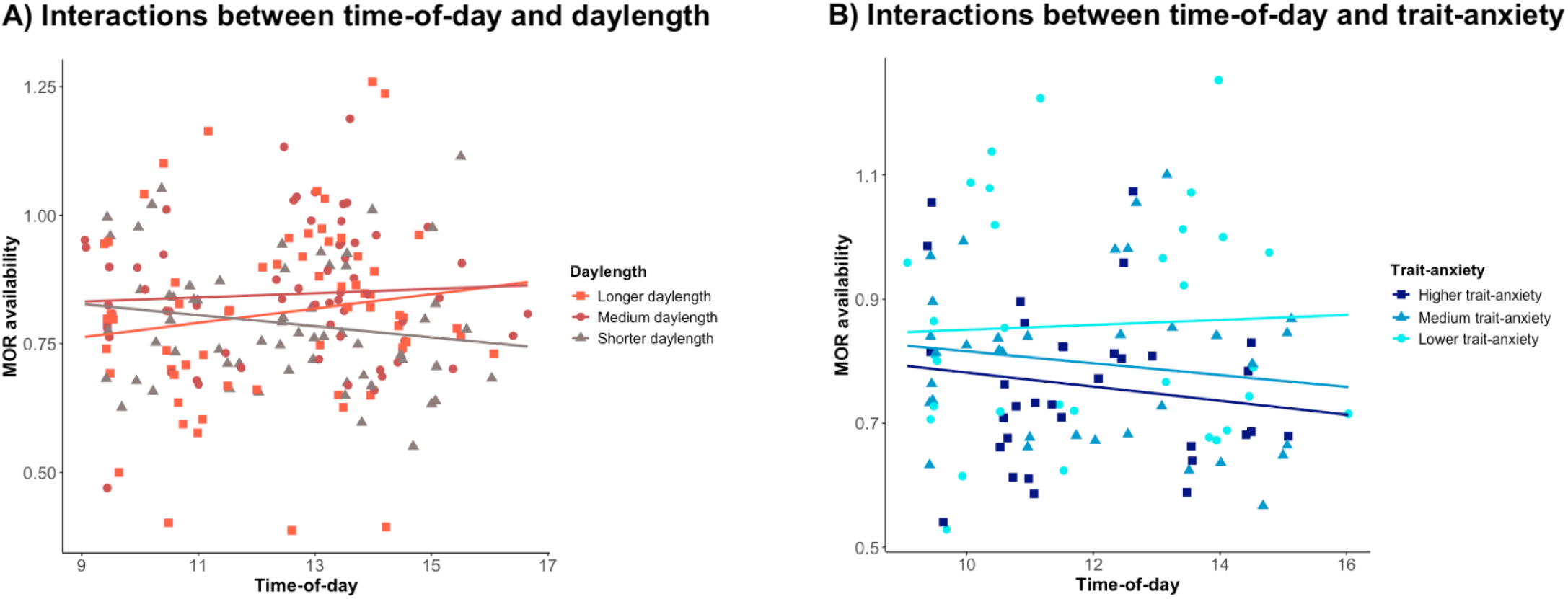
Diurnal patterns of mu-opioid receptor (MOR) availability is affected by daylength and trait-anxiety levels. The MOR availability data are the average values extracted from the significant clusters in each analysis, and plotted separately for daylength (**A**) and anxiety levels (**B**). Respective samples were stratified using a tertian split for visualisation purposes. The cut-off points for the daylength-related analysis were: shorter daylength < 9.5 h, medium daylength 9.5 h – 14.9 h, and longer daylength > 14.9 h. For the anxiety-related analysis, the following cut-off points were defined based on the State-Trait Anxiety Inventory (STAI-X; [Spielberger, 1989]) scores: lower trait-anxiety < 30, medium trait-anxiety 30 – 35, higher trait-anxiety > 35.

As for the interaction effects between trait-anxiety levels and daylength, the full volume model revealed no significant outcomes. However, the complementary ROI analysis yielded a nominally significant interaction between trait-anxiety and the second order polynomial of daylength in the right amygdala (T = -2.40; p_uncorr._ = 0.018; p_FDR_ = 0.511). Full data are presented in the **Supplementary Table S2**.

### 3.2. Analysis of D2R availability data

The analysis revealed diurnal variation in D2R signalling, showing that later TOD was associated with lower D2R availability in bilateral putamen and the left nucleus accumbens (p_FDR_ < 0.05; see **Table 1** and **Figure 4**). Trend-level decreases in D2R binding were also observed in bilateral caudate nucleus (p_FDR_ = 0.064). There were no interaction effects between TOD and daylength for any ROI.

**Table 1.** The effects of time-of-day and its interaction with daylength on the striatal D2R availability. Findings that survived the correction for multiple comparisons (p_FDR_ < 0.05) are presented in bold, while trend-level results (p_uncorr._ < 0.05) are additionally marked in italics.

| Region | Time-of-day (n = 184) | | | Time-of-day × daylength (n = 184) | | |
| --- | --- | --- | --- | --- | --- | --- |
| | t-stat | $\beta$ (SE) | $p_{\text{uncorr.}}$ ($p_{\text{FDR}}$) | t-stat | $\beta$ (SE) | $p_{\text{uncorr.}}$ ($p_{\text{FDR}}$) |
| L putamen | **-2.47** | **-0.00457**<br>(0.00185) | **0.015**<br>(0.039) | 0.14 | 0.00006<br>(0.00042) | 0.887<br>(0.942) |
| R putamen | **-2.47** | **-0.00475**<br>(0.00192) | **0.015**<br>(0.039) | 0.74 | 0.00032<br>(0.00435) | 0.460<br>(0.942) |
| L caudate nucleus | **-2.07** | **-0.00424**<br>(0.00205) | **0.040**<br>(0.064) | 0.29 | 0.00013<br>(0.00047) | 0.773<br>(0.942) |
| R caudate nucleus | **-2.16** | **-0.00449**<br>(0.00208) | **0.032**<br>(0.064) | 0.07 | 0.00003<br>(0.00047) | 0.942<br>(0.942) |
| L nucleus accumbens | **-2.53** | **-0.00566**<br>(0.00224) | **0.012**<br>(0.039) | -0.53 | -0.00027<br>(0.00051) | 0.595<br>(0.942) |
| R nucleus accumbens | -1.80 | -0.00399<br>(0.00222) | 0.073<br>(0.098) | -0.75 | -0.00038<br>(0.0005) | 0.453<br>(0.942) |
| L pallidum | -0.82 | -0.00179<br>(0.00217) | 0.411<br>(0.470) | 1.30 | 0.00063<br>(0.00049) | 0.197<br>(0.942) |
| R pallidum | -0.70 | -0.00155<br>(0.00221) | 0.484<br>(0.484) | 0.57 | 0.00028<br>(0.00050) | 0.571<br>(0.942) |

Abbreviations: SE, standard error; FDR, false discovery rate; L, left; R, right.

**Figure 4.**
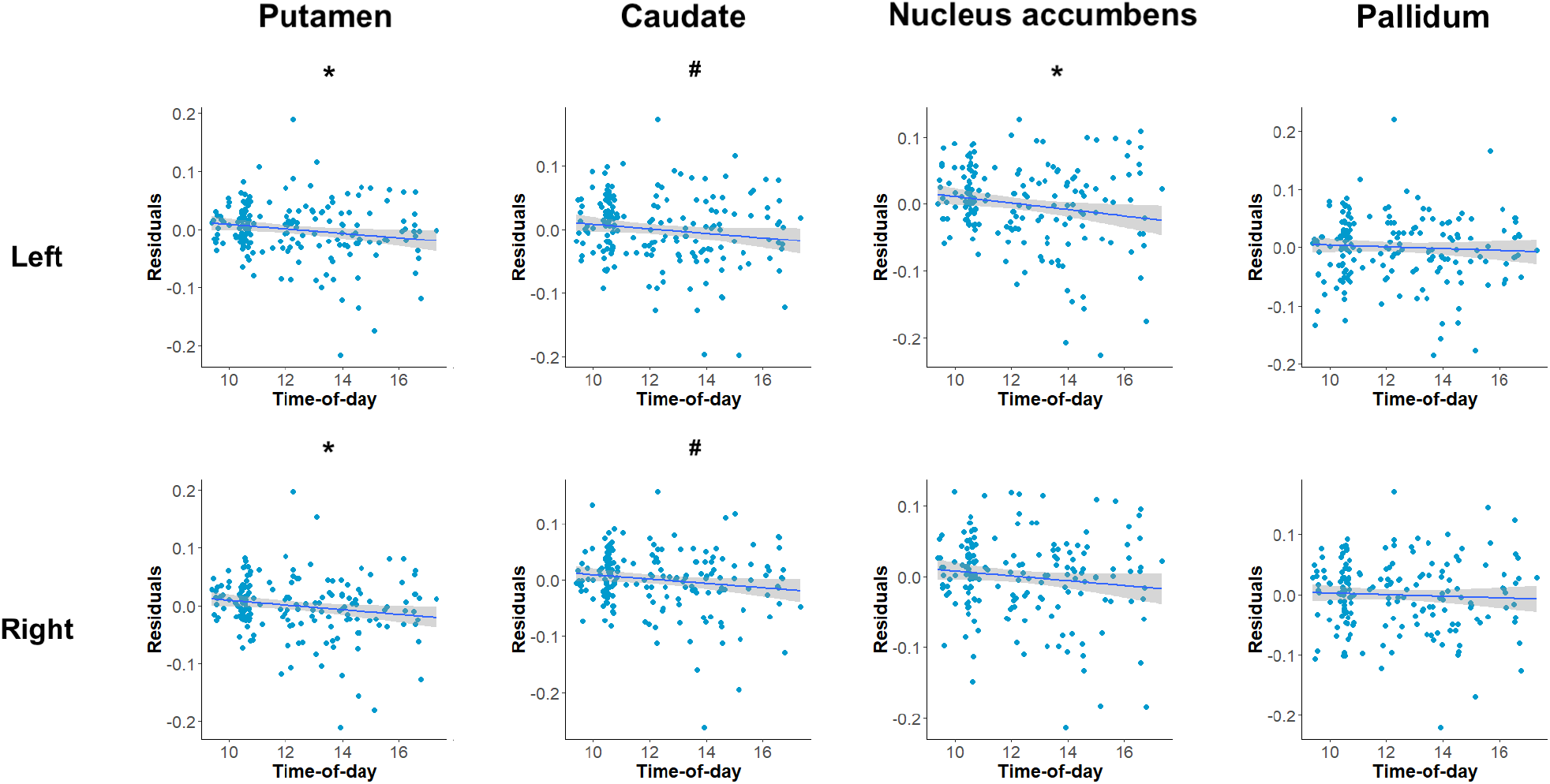
The effects of time-of-day (TOD) on D2R availability across the striatal regions. TOD was plotted against the residuals from the statistical models that excluded its main effect. Asterisks (*) denote statistically significant results after false discovery rate correction (p_FDR_ < 0.05), while hash (#) stands for significant findings before the multiple comparison correction (p_uncorr._ < 0.05).

## 4. Discussion

Our data show that both MOR and D2R availability demonstrate diurnal patterns, with later TOD scans having higher MOR and lower D2R availability levels. The seasonal daylength changes modulated the observed diurnal effects in MOR but not in D2R signaling. Specifically, in seasons with longer daylength, later TOD was associated with greater MOR availability, but this effect became less visible toward shorter days. In the meanwhile, the diurnal variation in MOR availability is also linked with levels of anxiety, showing that higher anxiety masks the main effect of TOD on MOR availability. Together, these findings highlight a unique role of the MOR system in mediating both types of rhythmic regulations (i.e., diurnal and seasonal rhythm), as well as anxiety traits. In contrast, the reduced D2R availability at later TOD and the independence of this effect from seasons highlights differences in diurnal processes between the two systems.

MORs are widely expressed across the human brain, with essential roles in social cognition and emotional processing [Hsu et al., 2013; Seppala et al., 2026; Zubieta et al., 2003]. The observed pattern of TOD variation within the limbic system may reflect diurnal changes in higher-order cognitive functions and mood, mirroring previous behavioural findings [Baranger et al., 2017; Marek et al., 2010]. Increased central MOR availability at later TOD also aligns with the reported reduction of peripheral endogenous opioid levels [Hindmarsh et al., 1989]. In addition, the probability of grooming-like behaviors, which stimulate the release of endogenous MOR agonists and are thought to buffer stress, is elevated in infants during the afternoon [Mai et al., 2026; Dunbar, 2018], consistent with the diurnal pattern observed in the present adult sample.

The diurnal change in MOR availability is further influenced by daylength and trait-anxiety levels. While the main effect of TOD on MOR availability is found more restricted to the frontal and inferior brain regions, its interaction effects with either daylength or anxiety levels span over substantially larger brain areas. Considerable topographic overlap underlying the two types of interaction effects further suggests that shorter daylength and higher anxiety levels may co-carry this modulatory effects on diurnal patterns of MOR signalling. Indirectly, this may further reflect an intimate relationship between dark seasons, anxiety, and MOR signalling. Our previous findings demonstrated seasonal variation in MOR signalling, but its behavioural relevance remained unclear [Sun et al., 2021]. In the present study, although interactions between daylength and behavioural traits were minimal, their similar modulation of TOD effects suggests a potential link between seasonal variation in MOR signalling and behaviour.

Potential mechanisms by which seasons and trait-anxiety levels modulate diurnal MOR trajectories remain to be explored. The seasonal driver effect may possibly reside in the altered length of night-time plateau of the melatonin level, which decreases during longer daylength [Cajochen and Schmidt, 2025]. Rodent data shows that the diurnal melatonin pattern influences the release of endogenous MOR agonists [Miguel Asai et al., 2007], either directly through the opioid-releasing neurons [Shavali et al., 2004], or indirectly through the stress-sensitive hypothalamus-pituitary-adrenal (HPA) axis [Yoshida et al., 2005; Klosen et al., 2019]. The HPA axis is also known to mediate the effects associated with trait-anxiety [Ariño-Braña et al., 2025]. Endogenous MOR agonists are released in response to stress, in turn regulating the HPA axis reactivity to stressors [Valentino and Van Bockstaele, 2015]. Chronic stress exposure leads to the development of opioid tolerance in rodents [Chaijale et al., 2013], and animals that develop anxiety-like behaviours due to chronic stress show decreased endogenous MOR signaling as compared to the resilient animals [Bérubé et al., 2014; Kavushansky et al., 2012]. In the meanwhile, we have shown that in rats which are subjected to seasonal light-dark cycling, the photoperiod-induced changes in corticosterone levels and striatal MOR availability are nominally correlated [Sun et al., 2021]. Therefore, the current finding may suggest a link between the stress-sensitive HPA functional axis, daylength and opioid signalling.

Our data also showed decreased striatal D2R availability at later TOD, in contrast to the previously reported lack of such differences between the morning and late evening [Earley et al., 2013]. Our finding may be partly explained by a theory that humans exhibit altered dopaminergic signalling in the middle of the active phase (i.e., midday for diurnal humans, midnight for nocturnal rodents) [Ferris et al., 2014], as driven by increased tonic dopamine levels [Ferris et al., 2014; Oskamp et al., 2017]. This may consequently lead to increased occupation of D2R by endogenous agonists, resulting in their lower availability at later TOD. D2R receptors are predominantly expressed in the striatum [Malén et al., 2022; Richfield et al., 1987], and elevated D2R signalling is often associated with higher perceived environmental reward rates and “wanting” responses [Bailey et al., 2020; Le Heron et al., 2020]. Previous studies have indeed reported elevated motivation levels of humans in the afternoon [Balter et al., 2024; Byrne et al., 2017b]. On the other hand, humans have also diminished reward learning [Whitton et al., 2018] and reduced striatal reactivity to monetary rewards during later TOD [Byrne et al, 2017a]. Therefore, this diurnal pattern of D2R signaling is supported, to some extent, by existing behavioural evidence.

### Limitations

The present work was based on historical brain PET data, where each participant was scanned only once to avoid a significant radiation load. Consequently, the findings reflect population-level patterns rather than within-subject variation. In a single PET scan, we cannot determine the specific molecular underpinnings but a combination of differences in receptor density, their affinity, or baseline occupancy by endogenous neurotransmitter. With regards to the diurnal processes, the available PET data was collected from the morning to the late afternoon, which prevented us from investigating potential curvilinear associations between receptor availability and TOD [Ferris et al., 2014]. This limited temporal coverage reflects the logistical complexity of PET imaging with ^11^C-labeled radiotracers, which depends on the operating hours of radiochemistry facilities. As for the seasonal effects, daylength was used as a seasonal regressor as in the previous studies [Sun et al., 2021; Sun et al., 2024], however, contributions from other seasonal factors, such as temperature, cannot be excluded. The seasonal modulation on diurnal processes should also be interpreted by taking into account the large magnitude of local photoperiodic variation, which may hinder the interpolation of the reported findings to regions with lower latitudes. Finally, given the correlational nature of this study, our data do not provide causal evidence regarding the effects of TOD, seasonal factors, or trait-anxiety.

## Conclusions

We conclude that both MOR- and D2R-related signalling in the human brain undergoes diurnal variations. The diurnal patterns observed in the MOR system are additionally modulated by seasonal factors and behavioural anxiety traits. The MOR system may therefore stand out as a potential therapeutic target for rhythm-tailored interventions in affective disorders, warranting future investigation. Furthermore, the present study establishes a baseline for diurnal variation of MOR signalling that may facilitate the identification of pathological alterations.

## 5. Statements and declarations

### 5.3. Author contributions

MRZ: Conceptualisation, Formal analysis, Writing - Original Draft, Visualisation, Funding acquisition. JR, JH, PN: Writing - Review & Editing. LN: Supervision, Writing - Review & Editing. LS: Conceptualisation, Supervision, Data Curation, Writing - Original Draft.

### 5.4. Data availability

The full-volume statistical maps generated during this study are available in the following NeuroVault collection: https://neurovault.org/collections/24436/. The Finnish legislation considers the medical imaging data as sensitive personal information that cannot be publicly shared, including in an anonymised format. Enquiries regarding the dataset can be sent by email to Lauri Nummenmaa or by post to Turku PET Centre c/o Turku University Hospital, Kiinamyllynkatu 4–8, FI-20520 Turku, Finland.

### 5.5. Ethics approval

Under the Finnish legislation, no ethical approval is required for register-based studies.

### 5.6. Consent to participate

The study does not require consents of participants as it is fully based on registered data.

