## Supplementary Material for "Diurnal and seasonal regulation of brain mu-opioid and dopamine D2 receptor signalling in humans"

**Supplementary Table S1.** The effects of time-of-day and its interactions with linear and quadratic daylength terms on the regional mu-opioid receptor (MOR) availability. Findings that survived the correction for multiple comparisons ($p_{FDR}$ < 0.05) are presented in bold, while trend-level findings ($p_{uncorr.}$ < 0.05) are additionally shown in italics.

| **Region** | **Time-of-day (n = 188)** | | | **Time-of-day × daylength (n = 188)** | | | **Time-of-day × daylength $^2$ (n = 188)** | | |
|---|---|---|---|---|---|---|---|---|---|
| | **t-stat** | **β (SE)** | **$p_{uncorr.}$ ($p_{FDR}$)** | **t-stat** | **β (SE)** | **$p_{uncorr.}$ ($p_{FDR}$)** | **t-stat** | **β (SE)** | **$p_{uncorr.}$ ($p_{FDR}$)** |
| L anterior cingulate cortex | 1.73 | 0.006<br>(0.004) | 0.086<br>(0.317) | **2.44** | **0.0015**<br>**(0.0006)** | **0.016**<br>**(0.034)** | -0.83 | -0.0001<br>(0.0001) | 0.406<br>(0.760) |
| R anterior cingulate cortex | 1.67 | 0.007<br>(0.004) | 0.097<br>(0.317) | **2.24** | **0.0015**<br>**(0.0007)** | **0.026**<br>**(0.047)** | -1.14 | -0.0001<br>(0.0001) | 0.256<br>(0.760) |
| L superior frontal gyrus | 1.46 | 0.006<br>(0.004) | 0.147<br>(0.374) | **2.86** | **0.0019**<br>**(0.0007)** | **0.005**<br>**(0.024)** | -0.78 | -0.0001<br>(0.0001) | 0.437<br>(0.760) |
| R superior frontal gyrus | ***2.06*** | ***0.008***<br>***(0.004)*** | ***0.041***<br>***(0.317)*** | **2.83** | **0.0020**<br>**(0.0007)** | **0.005**<br>**(0.024)** | -1.20 | -0.0001<br>(0.0001) | 0.231<br>(0.760) |
| L inferior frontal gyrus pars orbitalis | 0.79 | 0.003<br>(0.004) | 0.431<br>(0.500) | ***1.98*** | ***0.0014***<br>***(0.0007)*** | ***0.049***<br>***(0.070)*** | -0.32 | -0.0000<br>(0.0002) | 0.750<br>(0.886) |
| R inferior frontal gyrus pars orbitalis | 1.65 | 0.007<br>(0.004) | 0.102<br>(0.317) | **2.60** | **0.0019**<br>**(0.0007)** | **0.010**<br>**(0.029)** | -1.04 | -0.0002<br>(0.0002) | 0.301<br>(0.760) |
| L orbitofrontal cortex | 0.89 | 0.004<br>(0.004) | 0.375<br>(0.477) | **2.23** | **0.0016**<br>**(0.0007)** | **0.027**<br>**(0.047)** | -0.56 | -0.0001<br>(0.0002) | 0.580<br>(0.760) |
| R orbitofrontal cortex | 0.76 | 0.003<br>(0.004) | 0.446<br>(0.500) | **2.44** | **0.0017**<br>**(0.0007)** | **0.016**<br>**(0.034)** | -0.57 | -0.0001<br>(0.0002) | 0.569<br>(0.760) |
| L insula | 1.57 | 0.006<br>(0.004) | 0.119<br>(0.333) | **2.70** | **0.0017**<br>**(0.0006)** | **0.008**<br>**(0.029)** | -0.68 | -0.0001<br>(0.0001) | 0.496<br>(0.760) |
| R insula | 1.18 | 0.005<br>(0.004) | 0.238<br>(0.407) | **2.57** | **0.0017**<br>**(0.0006)** | **0.011**<br>**(0.029)** | -0.68 | -0.0001<br>(0.0001) | 0.500<br>(0.760) |
| L amygdala | ***2.35*** | ***0.008***<br>***(0.003)*** | ***0.020***<br>***(0.277)*** | 0.94 | 0.0005<br>(0.0005) | 0.348<br>(0.348) | -0.88 | -0.0001<br>(0.0001) | 0.379<br>(0.760) |
| R amygdala | ***3.11*** | ***0.010***<br>***(0.003)*** | ***0.002***<br>***(0.061)*** | 1.76 | 0.0010<br>(0.0006) | 0.080<br>(0.097) | -1.25 | -0.0001<br>(0.0001) | 0.215<br>(0.760) |
| L caudate nucleus | 1.31 | 0.006<br>(0.005) | 0.192<br>(0.407) | 1.46 | 0.0011<br>(0.0008) | 0.145<br>(0.162) | -0.27 | -0.0000<br>(0.0002) | 0.789<br>(0.886) |
| R caudate nucleus | 1.16 | 0.005<br>(0.004) | 0.246<br>(0.407) | 1.78 | 0.0013<br>(0.0008) | 0.076<br>(0.097) | -0.53 | -0.0001<br>(0.0002) | 0.595<br>(0.760) |
| L putamen | 1.83 | 0.006<br>(0.004) | 0.069<br>(0.317) | **2.60** | **0.0015**<br>**(0.0006)** | **0.010**<br>**(0.029)** | -0.27 | -0.0000<br>(0.0001) | 0.791<br>(0.886) |
| R putamen | ***2.02*** | ***0.007***<br>***(0.004)*** | ***0.045***<br>***(0.317)*** | **2.56** | **0.0015**<br>**(0.0006)** | **0.011**<br>**(0.029)** | -0.79 | -0.0001<br>(0.0001) | 0.430<br>(0.760) |
| L nucleus accumbens | 0.28 | 0.001<br>(0.004) | 0.777<br>(0.777) | 1.50 | 0.0009<br>(0.0006) | 0.136<br>(0.159) | 0.53 | 0.0001<br>(0.0001) | 0.597<br>(0.760) |

| | | | | | | | | | |
|---|---|---|---|---|---|---|---|---|---|
| R nucleus accumbens | 0.25 | 0.001<br>(0.003) | 0.251<br>(0.407) | 1.92 | 0.0011<br>(0.0006) | 0.056<br>(0.075) | 0.16 | 0.0000<br>(0.0001) | 0.873<br>(0.940) |
| L thalamus | 1.09 | 0.004<br>(0.004) | 0.276<br>(0.407) | ***2.16*** | ***0.0013***<br>***(0.0006)*** | ***0.032***<br>***(0.051)*** | -0.75 | -0.0001<br>(0.0001) | 0.456<br>(0.760) |
| R thalamus | 1.04 | 0.004<br>(0.004) | 0.298<br>(0.417) | ***2.08*** | ***0.0013***<br>***(0.0006)*** | ***0.039***<br>***(0.058)*** | -0.54 | -0.0001<br>(0.0001) | 0.588<br>(0.760) |
| L temporal pole | 1.35 | 0.006<br>(0.004) | 0.180<br>(0.407) | **2.26** | **0.0016**<br>**(0.0007)** | **0.025**<br>**(0.047)** | -0.69 | -0.0001<br>(0.0001) | 0.492<br>(0.760) |
| R temporal pole | 1.88 | 0.008<br>(0.004) | 0.062<br>(0.317) | ***2.18*** | ***0.0015***<br>***(0.0007)*** | ***0.031***<br>***(0.051)*** | -1.41 | -0.0002<br>(0.0001) | 0.161<br>(0.760) |
| L middle temporal gyrus | 0.93 | 0.004<br>(0.004) | 0.353<br>(0.471) | **3.40** | **0.0022**<br>**(0.0006)** | **<0.001**<br>**(0.001)** | -0.54 | -0.0001<br>(0.0001) | 0.591<br>(0.760) |
| R middle temporal gyrus | 0.80 | 0.003<br>(0.004) | 0.423<br>(0.500) | **3.10** | **0.0021**<br>**(0.0007)** | **0.002**<br>**(0.016)** | -0.86 | -0.0001<br>(0.0001) | 0.392<br>(0.760) |
| L inferior temporal gyrus | 1.15 | 0.004<br>(0.004) | 0.251<br>(0.407) | **3.33** | **0.0021**<br>**(0.0006)** | **0.001**<br>**(0.001)** | -0.74 | -0.0001<br>(0.0001) | 0.462<br>(0.760) |
| R inferior temporal gyrus | 1.11 | 0.004<br>(0.004) | 0.268<br>(0.407) | **3.45** | **0.0023**<br>**(0.0007)** | **<0.001**<br>**(0.001)** | -1.02 | -0.0001<br>(0.0001) | 0.311<br>(0.760) |
| L posterior cingulate cortex | -0.72 | -0.004<br>(0.005) | 0.471<br>(0.506) | 1.17 | 0.0010<br>(0.0008) | 0.244<br>(0.253) | 0.02 | 0.0000<br>(0.0002) | 0.988<br>(0.988) |
| R posterior cingulate cortex | -0.70 | -0.004<br>(0.005) | 0.488<br>(0.506) | 1.36 | 0.0012<br>(0.0009) | 0.176<br>(0.190) | -0.09 | -0.0000<br>(0.0002) | 0.927<br>(0.961) |

Abbreviations: SE, standard error; FDR, false discovery rate; L, left; R, right.

**Supplementary Table S2.** The effects of interactions of trait-anxiety with time-of-day, as well as the linear and quadratic daylength terms on the regional mu-opioid receptor (MOR) availability. Nominally significant associations are shown in bold and italics.

| Region | Trait-anxiety × time-of-day (n = 100) | | | Trait-anxiety × daylength (n = 100) | | | Trait-anxiety × daylength $^2$ (n = 100) | | |
|---|---|---|---|---|---|---|---|---|---|
| | t-stat | β (SE) | $p_{uncorr.}$ ($p_{FDR}$) | t-stat | β (SE) | $p_{uncorr.}$ ($p_{FDR}$) | t-stat | β (SE) | $p_{uncorr.}$ ($p_{FDR}$) |
| L anterior cingulate cortex | -1.78 | -0.0009 (0.0005) | 0.078 (0.192) | 0.89 | 0.00017 (0.00019) | 0.378 (0.928) | -0.94 | -0.00005 (0.00005) | 0.348 (0.872) |
| R anterior cingulate cortex | -1.84 | -0.0009 (0.0005) | 0.068 (0.192) | 0.33 | 0.00007 (0.00020) | 0.740 (0.928) | -1.03 | -0.00005 (0.00005) | 0.307 (0.872) |
| L superior frontal gyrus | -1.46 | -0.0008 (0.0005) | 0.148 (0.241) | 0.37 | 0.00008 (0.00021) | 0.714 (0.928) | -0.87 | -0.00005 (0.00006) | 0.388 (0.872) |
| R superior frontal gyrus | ***-2.01*** | ***-0.0011 (0.0005)*** | ***0.048 (0.192)*** | 0.23 | 0.00005 (0.00021) | 0.819 (0.928) | -0.69 | -0.00004 (0.00006) | 0.495 (0.872) |
| L inferior frontal gyrus pars orbitalis | ***-2.32*** | ***-0.0012 (0.0005)*** | ***0.023 (0.192)*** | 0.59 | 0.00013 (0.00021) | 0.558 (0.928) | -0.08 | -0.00000 (0.00006) | 0.935 (0.937) |
| R inferior frontal gyrus pars orbitalis | -1.50 | -0.0008 (0.0005) | 0.137 (0.241) | 0.30 | 0.00007 (0.00022) | 0.766 (0.928) | -0.26 | -0.00002 (0.00006) | 0.794 (0.937) |
| L orbitofrontal cortex | ***-2.53*** | ***-0.0012 (0.0005)*** | ***0.013 (0.183)*** | -0.63 | 0.00012 (0.00019) | 0.531 (0.928) | -0.66 | -0.00003 (0.00005) | 0.511 (0.872) |
| R orbitofrontal cortex | ***-3.03*** | ***-0.0014 (0.0005)*** | ***0.003 (0.090)*** | 1.03 | 0.00020 (0.00020) | 0.307 (0.928) | -0.70 | -0.00004 (0.00005) | 0.489 (0.872) |
| L insula | -1.95 | -0.0009 (0.0005) | 0.054 (0.192) | -0.17 | -0.00003 (0.00018) | 0.863 (0.928) | -0.47 | -0.00002 (0.00005) | 0.637 (0.872) |
| R insula | -1.76 | -0.0009 (0.0005) | 0.083 (0.192) | -0.29 | -0.00006 (0.00019) | 0.772 (0.928) | -0.87 | -0.00005 (0.00005) | 0.384 (0.872) |
| L amygdala | -1.20 | -0.0005 (0.0004) | 0.234 (0.345) | 0.33 | 0.00006 (0.00017) | 0.741 (0.928) | -1.83 | -0.00009 (0.00005) | 0.070 (0.818) |
| R amygdala | -0.93 | -0.0004 (0.0005) | 0.355 (0.414) | 0.88 | 0.00015 (0.00017) | 0.381 (0.928) | ***-2.40*** | ***-0.00011 (0.00005)*** | ***0.018 (0.511)*** |
| L caudate nucleus | -0.98 | -0.0006 (0.0006) | 0.331 (0.403) | -0.83 | -0.00020 (0.00025) | 0.409 (0.928) | -0.48 | -0.00003 (0.00007) | 0.629 (0.872) |
| R caudate nucleus | -0.67 | -0.0004 (0.0006) | 0.505 (0.544) | -1.04 | -0.00023 (0.00022) | 0.300 (0.928) | -0.45 | -0.00003 (0.00006) | 0.654 (0.872) |
| L putamen | -1.88 | -0.0008 (0.0004) | 0.063 (0.192) | -0.30 | -0.00005 (0.00018) | 0.766 (0.928) | -0.24 | -0.00001 (0.00005) | 0.814 (0.937) |
| R putamen | -1.72 | -0.0008 (0.0004) | 0.089 (0.192) | -0.75 | -0.00014 (0.00018) | 0.454 (0.928) | -0.69 | -0.00003 (0.00005) | 0.493 (0.872) |
| L nucleus accumbens | -1.10 | -0.0005 (0.0004) | 0.273 (0.349) | -1.01 | -0.00018 (0.00018) | 0.314 (0.928) | -0.14 | -0.00001 (0.00005) | 0.888 (0.937) |

| | | | | | | | | | |
|---|---|---|---|---|---|---|---|---|---|
| R nucleus accumbens | -0.01 | -0.0000<br>(0.0004) | 0.996<br>(0.996) | -0.98 | -0.00016<br>(0.00016) | 0.332<br>(0.928) | 0.12 | 0.00000<br>(0.00004) | 0.908<br>(0.937) |
| L thalamus | -0.73 | -0.0003<br>(0.0004) | 0.468<br>(0.524) | -0.78 | -0.00012<br>(0.00015) | 0.440<br>(0.928) | -0.48 | -0.00002<br>(0.00004) | 0.635<br>(0.872) |
| R thalamus | -0.62 | -0.0003<br>(0.0004) | 0.536<br>(0.556) | -0.21 | -0.00003<br>(0.00016) | 0.834<br>(0.928) | -1.47 | -0.00006<br>(0.00004) | 0.146<br>(0.818) |
| L temporal pole | ***-1.99*** | ***-0.0011***<br>***(0.0005)*** | ***0.049***<br>***(0.192)*** | 0.53 | 0.00011<br>(0.00021) | 0.599<br>(0.928) | -1.57 | -0.00009<br>(0.00006) | 0.121<br>(0.818) |
| R temporal pole | -1.43 | -0.0007<br>(0.0005) | 0.155<br>(0.241) | 0.33 | 0.00007<br>(0.00020) | 0.739<br>(0.928) | -1.57 | -0.00008<br>(0.00005) | 0.120<br>(0.818) |
| L middle temporal gyrus | -1.60 | -0.0008<br>(0.0005) | 0.113<br>(0.226) | -0.13 | -0.00003<br>(0.00021) | 0.895<br>(0.928) | -0.74 | -0.00004<br>(0.00006) | 0.463<br>(0.872) |
| R middle temporal gyrus | -1.47 | -0.0007<br>(0.0005) | 0.146<br>(0.241) | -0.29 | -0.00006<br>(0.00020) | 0.772<br>(0.928) | -0.47 | -0.00003<br>(0.00005) | 0.642<br>(0.872) |
| L inferior temporal gyrus | -1.80 | -0.0009<br>(0.0005) | 0.076<br>(0.192) | 0.48 | 0.00009<br>(0.00019) | 0.629<br>(0.928) | -1.34 | -0.00007<br>(0.00005) | 0.184<br>(0.859) |
| R inferior temporal gyrus | -1.76 | -0.0009<br>(0.0005) | 0.083<br>(0.192) | 0.14 | 0.00003<br>(0.00021) | 0.890<br>(0.928) | -1.03 | -0.00006<br>(0.00006) | 0.307<br>(0.872) |
| L posterior cingulate cortex | -1.13 | -0.0007<br>(0.0007) | 0.262<br>(0.349) | 0.09 | 0.00002<br>(0.00026) | 0.932<br>(0.932) | -0.16 | -0.00001<br>(0.00007) | 0.876<br>(0.937) |
| R posterior cingulate cortex | -1.10 | -0.0008<br>(0.0007) | 0.274<br>(0.349) | 0.31 | 0.00009<br>(0.00030) | 0.760<br>(0.928) | -0.08 | -0.00001<br>(0.00008) | 0.937<br>(0.937) |

Abbreviations: SE, standard error; FDR, false discovery rate; L, left; R, right.
